# Dopamine receptor 1 specific CRISPRa mice exhibit disrupted behaviors and striatal baseline cellular activity

**DOI:** 10.1101/2025.06.26.661721

**Authors:** Rianne R. Campbell, Mikah Green, Eric Y. Choi, Andreas B. Wulff, Allison N. Siclair, Smirti Khatri, Geralin Virata, Christina Barrett, Symphanie Key, Samir Patel, Mary Beth Rowell, Daniela Franco, Shanmugasundaram Ganapathy-Kanniappan, Brian N. Mathur, Mary Kay Lobo

## Abstract

The two main cell types in the striatum, dopamine receptor 1 and adenosine receptor 2a spiny projection neurons (D1-SPNs and A2A-SPNs), have distinct roles in regulating motor- and reward-related behaviors. Cre-selective CRISPR-dCas9 systems allow for cell-type specific, epigenomic-based manipulation of gene expression with gene-specific single guide RNAs (sgRNAs) and have potential to elucidate molecular mechanisms underlying striatal subtype mediated behaviors. Conditional transgenic Rosa26:LSL-dCas9-p300 mice were recently generated to allow for robust expression of dCas9-p300 expression with Cre-driven cell-type specificity. This system utilizes p300, a histone acetyltransferase which regulates gene expression by unwinding chromatin and making that region of the genome more accessible for transcription. Rosa26-LSL-dCas9-p300 mice were paired with Drd1-Cre and Ador2a-Cre mice to generate Drd1-Cre:dCas9-p300 and Ador2a-Cre:dCas9-p300 mouse lines and underwent behavioral phenotyping when sgRNAs were not present. Both Drd1-Cre:dCas9-p300 and Ador2a-Cre:dCas9-p300 have cell-type specific expression of spCas9 mRNA. Baseline behavioral assessments revealed that, under a sgRNA absent nontargeted state, Drd1-Cre:dCas9-p300 mice display repetitive spinning behavior, hyperlocomotion and enhanced acquisition of reward learning in comparison to all genotypic littermates. In contrast, Ador2a-Cre:dCas9-p300 do not exhibit any changes in behavior in comparison to their littermates. Electrophysiological recordings of dorsal striatum D1-SPNs revealed that Drd1-Cre:dCas9-p300 mice have increased input resistance and increased spontaneous excitatory postsynaptic current amplitude, together suggesting greater excitatory drive of D1-SPNs. Overall, these data demonstrate the necessity to validate CRISPR-dCas9 lines for research investigations. Additionally, the Drd1-Cre:dCas9-p300 line has the potential to be used to study underlying mechanisms of stereotypy and reward-learning.

**Significance Statement:** Using CRISPR-based tools to identify cell-type specific epigenomic and transcriptional mechanisms in disease and behavior has high utility for the neuroscience field. Previous limitations related to implementation of CRISPR-editing systems in mice were thought to be overcome by the generation of transgenic mouse lines, including a novel Cre-dependent dCas9-p300 mouse line. Our data shows however that Drd1-Cre:dCas9-p300 mice, generated from breeding Drd1-Cre mice with the dCas9-p300 mice, have cellular and behavioral disruptions under a nontargeted sgRNA absent state. Overall, these data suggest caution in employing CRISPR-dCas9 systems, particularly transgenic mouse lines, for research investigations.

## Introduction

Dysregulation of molecular processes within the brain can lead to the development of psychiatric disorders (Nestler and Lüscher, 2019). Changes in activity of the main projections within the striatum, spiny projections neurons (SPNs), are implicated in diseases related to reward and movement (Chandra et al., 2017; Fox and Lobo, 2019; Campbell and Lobo, 2023). SPNs can be further categorized into two different subtypes, based on gene expression differences, as Dopamine D1 receptor (D1R) expressing-SPNs and adenosine A2A receptors (A2A)-SPNs. Depending on the striatal subregion, these SPNs subtypes regulate opposing roles in a variety of behaviors (Canales and Graybiel, 2000; Lobo et al., 2010; Maia and Frank, 2011). Within the dorsal striatum (DSt), D1-SPN activity promotes motor behaviors, whereas A2A-SPN activity suppresses motor action. Imbalances in DSt SPN activity can lead to enhanced locomotion as well as stereotypies or repetitive, rhythmic behaviors, which are seen in disorders such as Tourette Syndrome and Autism Spectrum Disorders and Obsessive Compulsive Disorders (Albin and Mink, 2006; Guehl et al., 2008). Hyperactivation of D1-SPNs in mice induces stereotypies, including in dopamine-deficient mice (Chartoff et al., 2001; Lee et al., 2018). In the nucleus accumbens (NAc), D1-SPN activity promotes reward and motivated-related behaviors whereas A2A-SPN activity can often inhibit these behaviors (Lobo et al., 2010; Fox et al., 2023). Current research efforts are focused on identifying relevant changes in gene expression within that lead altered SPN activity and behaviors relevant to neuropsychiatric diseases.

The development of epigenome editing using CRISPR-Cas9 systems has allowed for precise interrogation of genes and molecular processes relevant to disease (Savell and Day, 2017; Savell et al., 2019; Choi et al., 2023). With nuclease-deactivated Cas9 (dCas9) fused with various transcription factors, gene editing tools have expanded to include chromatin modifiers, such as histone acetyltransferases, to promote or repress gene expression with the addition of a single guide RNA (sgRNA) which allows for precise targeting of a gene of interest. Recent studies have shown this system can be expanded to target multiple gene targets at once using multiplexed sgRNA constructs (Savell et al., 2020). When paired with Cre-induced expression, dCas9-fusion editors can epigenetically manipulate gene expression in a cell-type specific manner. However, dCas9-fusion editing systems have limitations, including often being too large for viral vector packaging (Asmamaw Mengstie, 2022). To overcome this and other challenges, Rosa26-LSL-dCas9-p300core mouse line were generated for in vivo epigenome editing for Cre-selective transcriptional activation of a gene of interest via sgRNA targeting (Gemberling et al., 2021). No studies to date have bred the novel Rosa26-LSL-dCas9-p300 mouse line with a Cre-expressing mouse line and assessed its behavioral validity for examining mechanisms relevant to neuropsychiatric diseases. Here, we generated Drd1-Cre:dCas9-p300 and Ador2a-Cre:dCas9-p300 mouse lines and tested their utility by characterizing baseline nontargeted-state differences of striatal-dependent behaviors between their littermates in the absence of sgRNAs.

## Materials and Methods

### Animal Subjects

Rosa26-LSL-dCas9-p300core (dCas9^p300^) mice (Gemberling et al., 2021), with an inserted transgene with a CAG promoter followed by a loxP-stop-loxP cassette with cDNA encoding Streptococcus pyogenes dCas9 fusion protein with p300 were obtained from the Jackson Laboratory. These mice were crossed with either D1-Cre (Line FK150) or A2A-Cre (Line KG139), both on a C57BL/6 background, to generate mice used for this study. Multiple breeding cages were used per cross, and breeders were refreshed every 9 months. All pups from these crosses were genotyped and all genotypes (Drd1-Cre:dCas9-p300, Ador2a-Cre:dCas9-p300, dCas9-p300, Drd1-Cre, Ador2a-Cre, wildtype) were used for behavioral studies. We observed similar numbers of male and female offspring within each cross. All behavioral and molecular studies were conducted with male and female mice were between 2-5 months old and group housed for all experiments. Mice were given food and water ad libitum and housed in University of Maryland School of Medicine animal facilities on a 12-hour light/dark cycle. All experiments were performed in accordance with the Institutional Animal Care and Use Committee guidelines at the University of Maryland School of Medicine.

### Behavioral Tests

All animals underwent behavioral experiments in the following order to minimize cohort-effects and reduce carryover anxiety from prior testing. Mice were initially weighed and then underwent open field testing and followed by elevated plus maze testing 24 hours later. After a minimum of 48 hours off, all mice underwent food self-administration testing. 24 hours after the last food self-administration testing, mice from the D1-Cre x dCas9-p300 crosses underwent spinning tests.

### Open Field Test (OFT)

To assess baseline locomotor activity and anxiety-like behaviors, mice were placed in 43cm x 43cm white box acrylic box and allowed to explore the arena for 15 minutes. Amount of time spent in the corners versus centers of the arena was tracked along with total distance traveled in arena.

### Elevated Plus Maze (EPM)

To evaluate anxiety-like behaviors, mice were placed in the center of an elevated plus maze apparatus and behaviors were recorded for 5 minutes. The percent of time spent in the open versus closed arms was recorded for 5 minutes. Video-tracking was conducted with TopScan Lite software (Cleversys, Reston, VA, USA) for both OFT and EPM in similar manner to previous studies (Fox et al., 2023; Morais-Silva et al., 2023).

### Spinning Test

For the spinning test, mice were placed in a novel, clean cage and number of full 360° spins were counted for 5 minutes by a blinded experimenter, similar to previous studies (Engeln et al., 2021).

### Food Self-administration

To examine baseline reward learning, mice were trained to nose poke in an operant chamber (Med Associates) for food pellets (Bio-Serv: #F05684) paired with a 10 second illumination of light cue under a fixed-ratio 1, followed by a 10 second time out (FR1TO10 s) for 2 hours daily for 10 consecutive days. During the time out period, nose pokes in the active port were recorded but did not trigger additional pellet delivery. Throughout the session, responses on the inactive nose poke were recorded but had no programmed consequences.

### RNAScope

Brains were rapidly extracted and flash frozen for 20s in isopentane at −40°C and then stored at −80°C until later use. 16 micrometer coronal sections containing the striatum were collected in a −20°C cryostat (Leica), mounted on Superfrost Plus slides (Fisher Scientific) and stored at −80°C until the day of RNAScope assay. We used the RNAScope Multiple Fluorescence Reagent V1 Kit (Advanced Cell Diagnostics) according to the user manual for fresh-frozen tissue and as previously described (Maynard et al., 2018). Briefly, tissue sections were submerged in a 10% neutral buffered formalin solution (Cat # HT501128 Sigma-Aldrich, St. Louis, Missouri) for 20 min fixation followed by protease IV pretreatment for 20 min. Sections were then incubated with a custom-designed Channel 3 *Cas9*-C3 probe (519411-C3) and a commercially available *Drd1* probe (Cat # 461901Advanced Cell Diagnostics, Hayward, California) and *Ador2a* probe (Cat # 409431-C2 Advanced Cell Diagnostics, Hayward, California). Probes were fluorescently labeled with orange (excitation 550 nm), green (excitation 488 nm), or far red (excitation 647) fluorophores using the Amp 4 Alt B-FL. Confocal images were acquired in at 40x and 63x magnification using a Lecia Sp8 confocal microscope.

### RNAScope Analysis

For analysis, number of Cas9 puncta in Drd1 versus Ador2a cells were counted using Imaris software. Total number of Cas9 positive Drd1 cells or Ador2a cells out of the total number of Drd1 or Adora2a cells was quantified in each slice per mouse (three - four slices per mouse). Additionally, the average number of Cas9+ cells per cell type in each subject was then used to calculate the percentage of total positive dCas9 Drd1 or Adora2a cells.

### RNA Extraction and qRT-PCR

Two 14-gauge nucleus accumbens punches and two 12-gauge dorsal striatum per mouse were collected 24 hours after the last behavior and stored at −80°C until processing. RNA was extracted as previously described (Franco et al., 2022). Samples were dounce homogenized using TRIzol (Invitrogen; #15596018) and extracted following chloroform-based phase separation and centrifugation. Samples were then further purified using the EZNA MicroElute Total RNA kit (Omega Bio-Tek, Norcross, GA, United States; #R6831-01) with a DNA denaturing DNase step (Qiagen, Germantown, MD, United States; #79254). RNA concentration and quality were determined with a NanoDrop 1000 spectrophotometer (Thermo Fisher Scientific). 400 ng of complementary DNA (cDNA) was synthesized using the reverse transcriptase iScript complementary DNA synthesis kit (Bio-Rad, Hercules, CA, United States; # 1708891), then diluted to a final concentration of 2 ng/μL. Relative mRNA expression changes were measured by quantitative PCR using Perfecta SYBR Green FastMix (Quantabio, Beverly, MA, United States; #95072) with a Bio-Rad CFX384 qPCR system. Quantification of mRNA changes was performed using the −ΔΔC_T_ method described previously (Chandra et al., 2015). List of primers: *Drd1:* Forward: GAGCGTGGTCTCCCAGAT; Reverse: GGATGCTGCCTCTTCTTCTG; *Ador2a* Forward: CACGCAGAGTTCCATCTTCA; Reverse: AATGACAGCACCCAGCAAAT; p300 Forward: CTTCAGACAAGTCTTGGCATAGT; Reverse: CCGACTCCCATGTTGAGGTTT; *GAPDH:* Forward: AGGTCGGTGTGAACGGATTTG; Reverse: TGTAGACCATGTAGTTGAGGTCA.

### Western Blotting

Western blot procedures were performed as described previously (Chandra et al., 2015; Choi et al., 2023). Frozen striatal punches were syringe homogenized in 30 μl of lysis buffer containing 320 mM sucrose, 5 nM HEPES buffer, 1% SDS, phosphatase inhibitor cocktails I and II (Sigma) and protease inhibitors (Roche), followed by sonication using an ultrasonic processor (Cole Parmer). Protein concentrations were determined using the BCA protein assay (Thermo Fisher Scientific) and 15–20 μg samples of total protein were loaded onto 10 well Mini-Protean TGX Gels (Bio-Rad 4569033). Samples were transferred onto a nitrocellulose membrane (Bio-Rad 1620094) and blocked for 1 h in blocking buffer (5% nonfat dry milk in Tris-buffered saline, pH 7.6, with 0.1% Tween). Blocked membranes were incubated overnight at 4°C in blocking buffer with primary antibodies (mouse anti-SpCas9 (1:500; Encor MCA-3FP), Rabbit anti-GAPDH (1:2000; CST: 2118S)). Membranes were then incubated with horseradish peroxidase-labeled secondary antibodies (Vector Laboratories, catalog #PI-1000, 1:20,000) in blocking buffer. The bands were visualized using SuperSignal West Dura Extended Duration substrate (Pierce, catalog #34075). Bands were quantified with Image Lab Software (Bio-Rad) and normalized to GAPDH to control for equal loading.

### Acute slice preparation

Male and female Drd1-Cre and Drd1-Cre:dCas9-p300 mice (2–3 months) were deeply anesthetized with isoflurane before rapid decapitation and brain removal. 250 µm coronal sections were collected in ice-cold modified artificial cerebral spinal fluid (aCSF: 95% oxygen, 5% carbon dioxide, 194 mM sucrose, 30 mM NaCl, 4.5 mM KCl, 1 mM MgCl2, 26 mM NaHCO3, 1.2 mM NaH2PO4, and 10 mM D-glucose) using a Leica VT1200 vibrating microtome. Brain sections were then transferred to regular aCSF (95% oxygen, 5% carbon dioxide, 124 mM NaCl, 4.5 mM KCl, 2 mM CaCl2, 1 mM MgCl2, 26 mM NaHCO3, 1.2 mM NaH2PO4, and 10 mM D-glucose) and incubated at 32°C for 30 min before being stored at room temperature until recording.

### Whole-cell patch clamp electrophysiology

Slices were hemisected, placed into a recording chamber, and perfused with temperature controlled aCSF (29–31 °C) during the recording procedure. Dorsal striatum D1R, mCherry-positive medium spiny neurons were fluorescently visualized. Borosilicate glass pipettes (resistances ranging from 3–5MΩ) were pulled using a Narishige PC-100 micropipette puller and used for whole-cell patch clamp recordings. All recordings were amplified with a MultiClamp 700B amplifier (Molecular Devices), filtered at 2 kHz, digitized at 10 kHz and acquired using Clampex 10.4.1.4 software (Molecular Devices).

For recordings performed in current clamp configuration, recording pipettes were filled with a potassium-based internal solution (300–305 mOsm; pH 7.3) composed of (in mM) 126 potassium gluconate, 4 KCl, 10 HEPES, 4 ATP-Mg, 0.3 GTP-Na and 10 phosphocreatine. Membrane capacitance and resistance were collected at –60 mV in voltage clamp configuration, whereas the resting membrane potential (RMP) was obtained in current clamp configuration. Cells were permitted to equilibrate with the recording pipette internal solution for 5 minutes prior to data collection. Intrinsic excitability was assessed using a series of depolarizing current injections (0–460 pA; 40 pA step at 500 ms each). Recordings were analyzed for action potential (AP) frequency, threshold and latency. Rheobase (pA) was measured by ramping up current injection over 800 ms (max current 400-600 pA) and input resistance was measured as the slope of the depolarization induced by the ramped current injection. Current clamp data were analyzed in MATLAB (R2021A).

For recordings performed in voltage clamp configuration, recording pipettes were filled with a cesium-based internal solution (297-300 mOsm; pH 7.3) composed of (in mM) 120 CsMeSO3, 10 HEPES, 5 NaCl, 10 TEA-Cl, 1.1 EGTA, 0.3 Na-GTP and 4 Mg-ATP. Spontaneous excitatory postsynaptic currents (sEPSCs) were isolated with the addition of 50μM picrotoxin to the aCSF. Five minute recording epochs were collected for sEPSC amplitude (pA) and inter-event-interval (ms) analysis. Voltage clamp data were analyzed offline in MiniAnalysis (Synaptosoft).

### Statistical Analysis

Data are presented as mean ±SEM with individual data points overlaid. All statistical tests were performed in Graph Pad Prism 10. Normality was assessed with Shapiro-Wilk test. One-way ANOVAs were run for DSt qPCR (Drd1-dCas9-p300 lines: Ador2a, Drd1, p300; Ador2a-dCas9-p300 lines: Drd1, p300) and for NAc qPCR (Drd1-dCas9-p300 lines: p300; Ador2a-dCas9-p300 lines: Adora2a) and Kruskal-Wallis tests were used for DSt qPCR (Ador2a-dCas9-p300 lines: Ador2a) and NAc qPCR (Drd1-dCas9-p300 lines: Ador2a, Drd1; Ador2a-dCas9-p300 lines: Drd1, p300). One-way ANOVAs were used for baseline behavioral assessments (Weight, stereotypy, locomotion, OFT center %) when comparing across genotype. When reviewing behavioral assessments with sex as a variable, two-way ANOVAs were used for EPM, and food self-administration in addition to assessing differences in weight, stereotypy, locomotion, OFT center %, and for EPM with sex as a variable. Three-way ANOVA was used for comparisons of food self-administration between Drd1-Cre:dCas9-p300 genotypes and sex. Tukey post-hoc tests were used for parametric tests and Dunn’s multiple testing for non-parametric tests. The following tests were used to assess changes in whole-cell electrophysiology measurements: unpaired t-tests were used for: input resistance, rheobase, threshold potential, and sEPSC inter-event intervals; unpaired t-test with welch’s correction was used for sEPSC amplitude; a two-way ANOVA was conducted for AP potential frequency. Samples were excluded if they failed Grubbs’ outlier test. Sample sizes were determined from previous studies using mouse behavior, qRT-PCR, western blotting and electrophysiology with mouse tissues (Chandra et al., 2015; Engeln et al., 2021; Fox et al., 2023).

## Results

### Cell-type selective Cre-dependent Cas9 expression in the striatum within Drd1-Cre:dCas9-p300 and Ador2a-Cre:dCas9-p300 mice

Heterozygous conditional transgenic Rosa26:LSL-dCas9-p300 mice were paired with hemizygous Drd1-Cre and hemizygous Ador2a-Cre mice to generate mice that would allow for use of the dCas9-p300 fusion system, or CRISPR activation system, in a SPN subtype-specific manner (**Fig.1A**). Drd1-Cre pairings results in the following genotypes within their offspring: Drd1-Cre^+/-^::dCas9-p300^+^ (Drd1-Cre:dCas9-p300), Drd1-Cre^-/-^::dCas9-p300^+^(dCas9-p300), Drd1-Cre^+/-^::dCas9-p300^-^(Drd1-Cre), and Drd1-Cre^-/-^::dCas9-p300^-^(wildtype). Similarly, Ador2a-Cre breeding results in the following genotypes within their offspring: Ador2a-Cre^+/-^::dCas9-p300^+^ (Ador2a-Cre:dCas9-p300), Ador2a ^-/-^::dCas9-p300^+^(dCas9-p300), Ador2a -Cre^+/-^::dCas9-p300^-^(Ador2a-Cre), and Ador2a-Cre^-/-^::dCas9-p300^-^ (wildtype). To confirm the cell-type specificity of Cas9-p300 expression in Drd1-Cre:dCas9-p300 and Ador2a-Cre:dCas9-p300 mice, fluorescence in situ hybridization was performed to examine expression of Cas9, Drd1 and Ador2a in the striatum (**Fig. 1B, Table S1**). Within Drd1-Cre:dCas9-p300 mice, Drd1 expressing (Drd1+) cells have significantly higher Cas9 mRNA expression in comparison to Ador2a positive (Ador2a+) cells (unpaired t-test; t(4)=8.096, p=0.0013). In the Ador2a-Cre:dCas9-p300 mice, Ador2a+ cells have significantly higher Cas9 mRNA expression in comparison to Drd1+ cells in the NAc (unpaired t-test; t(4)=6.069, p=0.0037). SpCas9 protein in the striatum was observed only in mice with Cre and dCas9-p300 transgene expression (**Fig. 1C, S1**). These data demonstrate that Drd1-Cre:dCas9-p300 and Ador2a-Cre:dCas9-p300 mice have Cre-dependent SPN subtype-specific expression of the dCas9-p300 gene in the striatum.

**Figure 1.**
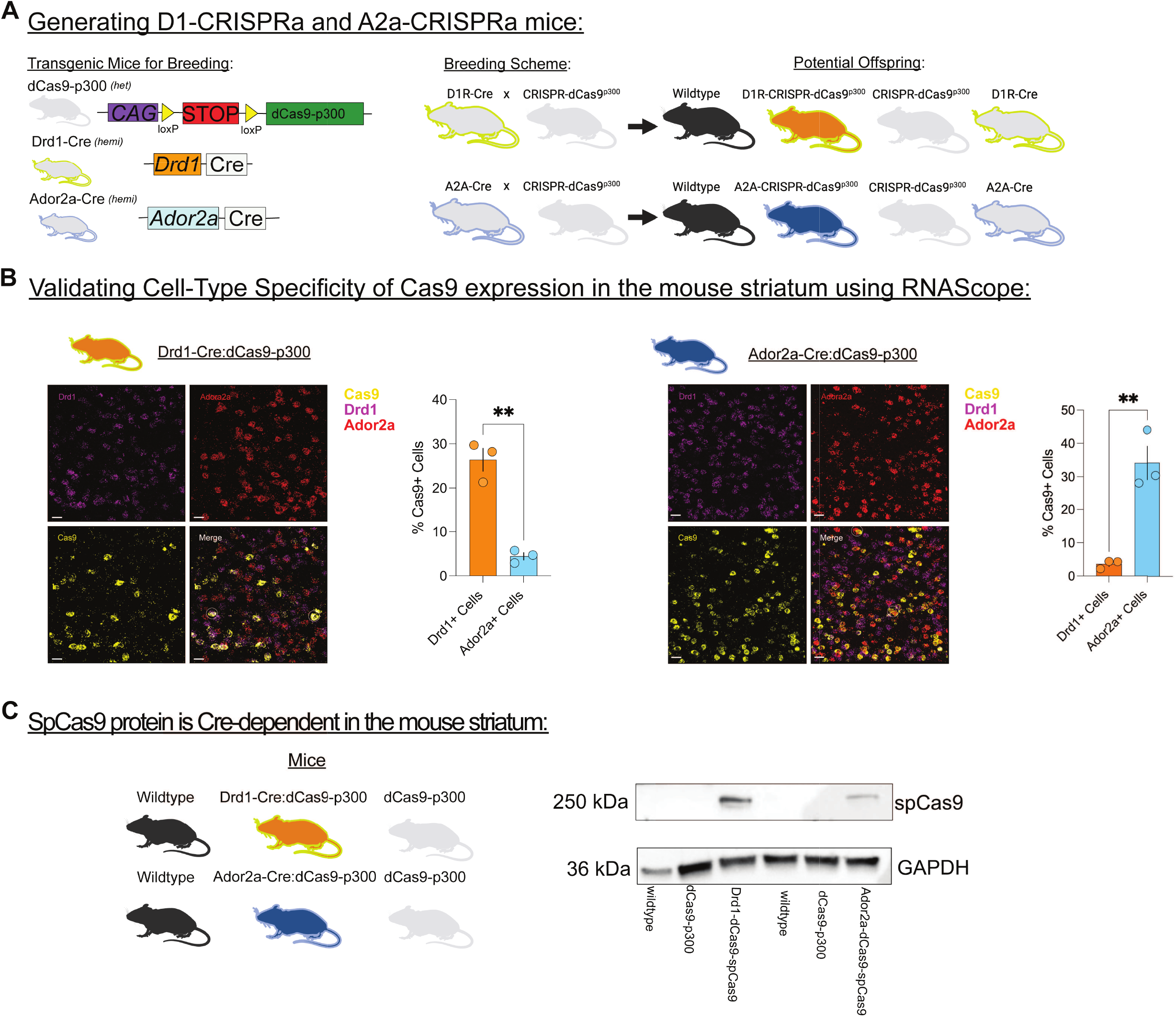
Cre-dependent Cas9 expression in the striatum within Drd1-Cre: and Ador2a-Cre:dCas9-p300 mice. **A**, The conditional transgenic Rosa26:LSL-dCas9-p300 mice were paired with Drd1-Cre mice which resulted in the following potential offspring: Drd1-Cre:dCas9-p300, dCas9-p300, Drd1-Cre, and wildtype littermates. Similarly, Ador2a-Cre mice were bred the conditional transgenic Rosa26:LSL-dCas9-p300 mice which can result in the following potential offspring: Ador2a-Cre:dCas9-p300, dCas9-p300, Ador2a-Cre, and wildtype littermates. **B**, (left) Drd1 expressing (Drd1+) cells have significantly high Cas9 mRNA expression in comparison to Ador2a positive (Ador2a+) cells in the nucleus accumbens (NAc) of Drd1-Cre:dCas9-p300 mice (n=3 mice). (right) Ador2a+ cells have significantly high Cas9 mRNA expression in comparison to Drd1+ cells in the NAc of Ador2a-Cre:dCas9-p300 mice (n=3 mice). Representative cells with overlapping expression for Drd1 or Ador2a and Cas9 are circled in white. In the image panel each scale bar is 20μm. Each bar graph represents mean ±SEM. *p<0.05, **p<0.01. **C**, Western blotting was performed from striatum from Drd1-Cre:dCas9-p300 and Ador2a-Cre:dCas9-p300 mice as well as their wildtype and dCas9-p300 littermates. Expression of the spCAS9 protein was detected in Drd1-Cre:dCas9-p300 and Ador2a-Cre:dCas9-p300 samples, but not their wildtype or dCas9-p300 counterparts. GAPDH however was detected in all samples.

### Genotypic differences in expression levels of Drd1, Ador2a and p300 in the striatum from cell-type specific dCas9-p300 transgene expression

To determine whether transgene dCas9-p300 expression impacts baseline mRNA expression in the striatum, RT-qPCR was performed on NAc and DSt tissues and compared across all genotypes related to Drd1-Cre:dCas9-p300 (**Fig. 2A, B**) and Ador2a-Cre:dCas9-p300 lines (**Fig. 2C, D**). NAc expression levels of *Drd1* (Kruskal-Wallis: H_(3)_=4.619, p=0.2020), *Ador2a* (Kruskal-Wallis: H_(3)_=3.711, p=0.2944) and p300 (One-way ANOVA: F_(3, 37)_=1.205, p=0.3213) did not differ between Drd1Cre:dCas9-p300, dCas9-p300, Drd1-Cre, and wildtype mice. Similarly, no changes in DSt expression levels of *Drd1* (One-way ANOVA: F_(3,41)_=1.570, p=0.2112) or *Ador2a* (one-way ANOVA: F_(3,41)_=1.014, p=0.3965) were seen between genotypes. However, Drd1-Cre:dCas9-p300 had significantly higher expression levels of DSt *p300* mRNA in comparison to dCas9-p300 mice (one-way ANOVA: F_(3,41)_=3.103, p=0.0369). Ador2a-Cre:dCas9-p300, dCas9-p300, Ador2a-Cre, and wildtype mice also had similar expression levels of *Drd1* (Kruskal-Wallis: H_(3)_=6.140, p=0.1050), *Ador2a* (one-way ANOVA: F_(3,24)_=1.674, p=0.1990) and p300 (Kruskal-Wallis: H_(3)_=2.898, p=0.4077) mRNA in the NAc. Similarly, no changes in DSt expression levels of p300 (one-way ANOVA: F_(3,24)_=0.3522, p=0.7879) were seen between Ador2a-Cre:dCas9-p300, dCas9-p300, Ador2a-Cre, and wildtype mice. Reductions in *Ador2a* expression within DSt of Ador2a-Cre:dCas9-p300 in comparison to wildtypes was observed (Kruskal Wallis: H_(3)_=13.10, p=0.0044: *post-hoc*: wildtype versus Ador2a-Cre:dCas9-p300: p=0.0026). Reductions in *Drd1* expression within Ador2a-Cre DSt samples in comparison *to* Ador2a-Cre:dCas9-p300 and dCas9-p300 were also detected (one-way ANOVA: F_(3,24)_=4.960, p=0.0081; Tukey’s *post hoc*: Ador2a-Cre:dCas9-p300 vs. Ador2a-Cre: p<0.05; dCas9-p300 vs. Ador2a-Cre: p<0.05).

**Figure 2.**
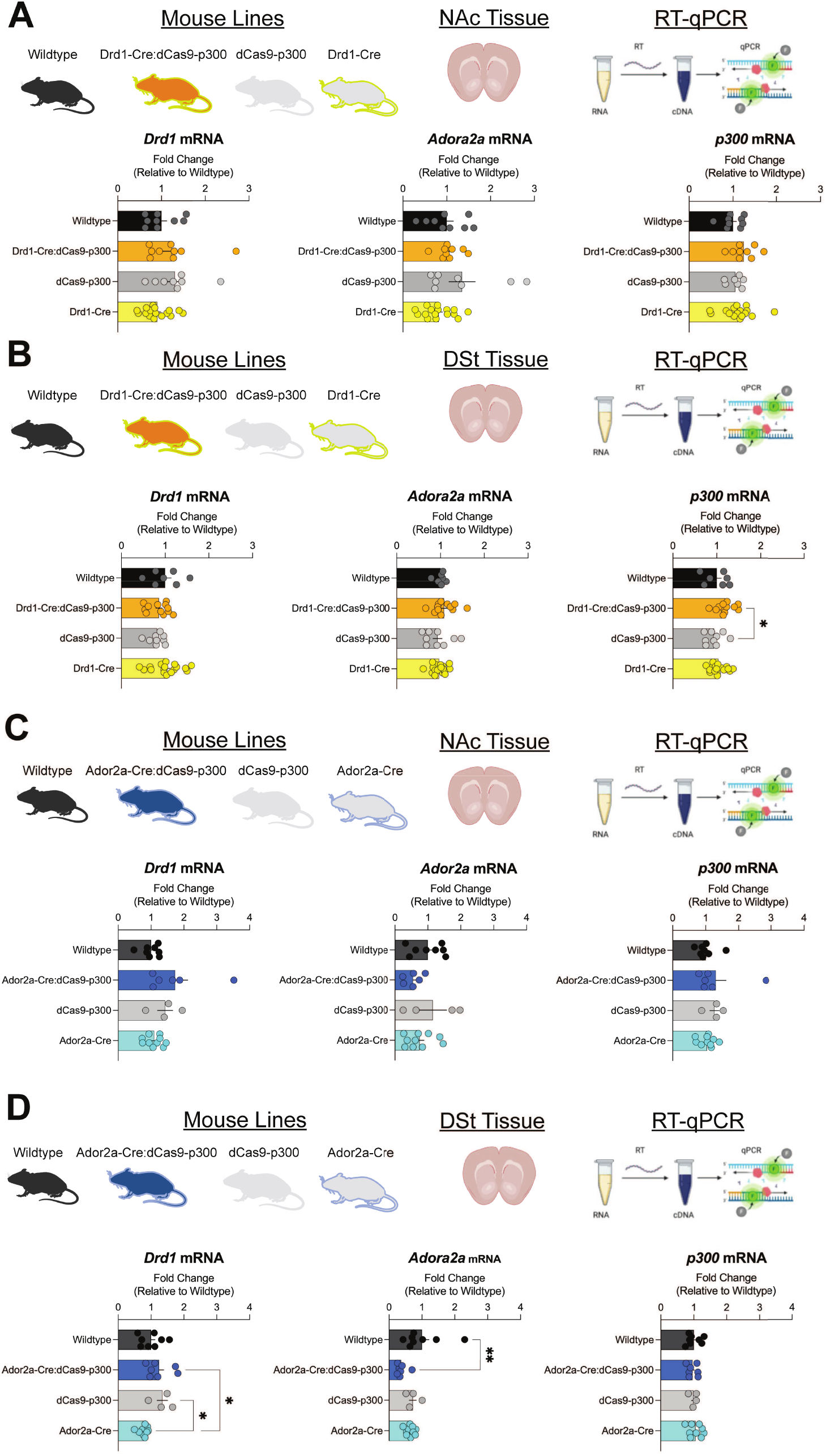
Expression levels of Drd1, Ador2a and p300 in the striatum are not altered by cell-type specific dCas9-p300 transgene expression. **A**, Nucleus accumbens (NAc) tissue from Drd1-Cre:dCas9-p300, dCas9-p300, Drd1-Cre, and wildtype mice was collected and processed for RT-qPCR (samples per genotype: n=8-16). Expression levels of *Drd1* and *Ador2a*, did not differ between genotypes. p300 expression is higher in NAc Drd1-Cre:dCas9-p300 samples in comparison to dCas9-p300 samples. **B**, Dorsal Striatum (DSt) tissue from Drd1-Cre:dCas9-p300, dCas9-p300, Drd1-Cre, and wildtype mice was collected and processed for RT-qPCR (samples per genotype: n=7-17). Expression levels of *Drd1, Ador2a*, and p300 did not differ between genotypes. **C.** NAc tissue from Ador2a-Cre:dCas9-p300, dCas9-p300, Ador2a-Cre, and wildtype mice was collected and processed for RT-qPCR(samples per genotype: n=4-10). Expression levels of *Drd1, Ador2a*, and p300 did not differ between genotypes. **D**. DSt from Ador2a-Cre:dCas9-p300, dCas9 -p300, Ador2a-Cre, and wildtype mice was collected and processed for RT-qPCR (samples per genotype: n=4-10). Expression levels of p300 were unchanged across genotypes. DSt *Drd1* expression is reduced in Ador2a-Cre samples in comparison to dCas9-p300 and Ador2a-Cre:dCas9-p300 DSt samples. Ador2a-Cre:dCas9-p300 DSt samples had reduced expression of *Ador2a* in comparison of DSt wildtype samples. Each bar represents mean +SEM. *p<0.05, **p<0.01.

### Drd1-Cre:dCas9-p300 mice have augmented baseline locomotion, stereotypic spinning and reward-learning behaviors

#### Adult Baseline Weight and Activity

The effects of Cas9-p300 expression in Drd1 cells on bodyweight and home-cage behaviors were first assessed on Drd1-Cre:dCas9-p300 mice and their littermates. dCas9-p300, but not Drd1-Cre:dCas9-p300 mice exhibit increased weight in comparison to other genotypes (**Fig. 3A**: one-way ANOVA; F_(3,57)_=3.493, p=0.0213; Tukey’s *post-hoc*: wildtype vs. dCas9-p300: p<0.05). However, when examining with separate sex, no genotypic differences were observed. As expected, male mice had significantly higher body weights than females (**Fig. 3B:** Two-way ANOVA: Interaction: F_(3,63)_=0.64, p=0.5892; Sex: F_(1,63)_=16.35, p=0.0001; Genotype: F_(3,63)_=1.453, p=0.2360). No notable differences in grooming or appearance were observed. However stereotypic locomotor behavior was observed in Drd1-Cre:dCas9-p300 mice. Baseline differences in locomotor activity in an open field were also examined. Drd1-Cre:dCas9-p300 mice had significantly higher locomotion in comparison to wildtype, dCas9-300, and Drd1-Cre mice (**Fig. 3C:** one-way ANOVA F_(3,65)_=7.753, p<0.001; Tukey’s *post-hoc*: wildtype vs Drd1-Cre:dCas9-p300: p<0.01; Drd1-Cre:dCas9-p300 vs dCas9-p300: p<0.001; Drd1-Cre:dCas9-p300 vs Drd1-Cre: p<0.001;). Male Drd1-Cre:dCas9-p300 mice have significantly higher locomotion than female Drd1-Cre:dCas9-p300, as well as all other male and female genotypes (**Fig. 3D:** two-way ANOVA: Interaction: F_(3,70)_=7.01, p<0.003; Sex: F_(1,70)_=7.09, **p=0.001; Genotype:F_(3,70)_=8.01, p=0.0001; Sidak’s *post-hoc*: Drd1-Cre:dCas9-p300 vs. all other genotypes p<0.01). Together, these results would suggest disrupted locomotor stereotypy and hyperlocomotion may occur from functional promiscuity of canonical dCas9-p300 expression in Drd1-expressing cells.

**Figure 3.**
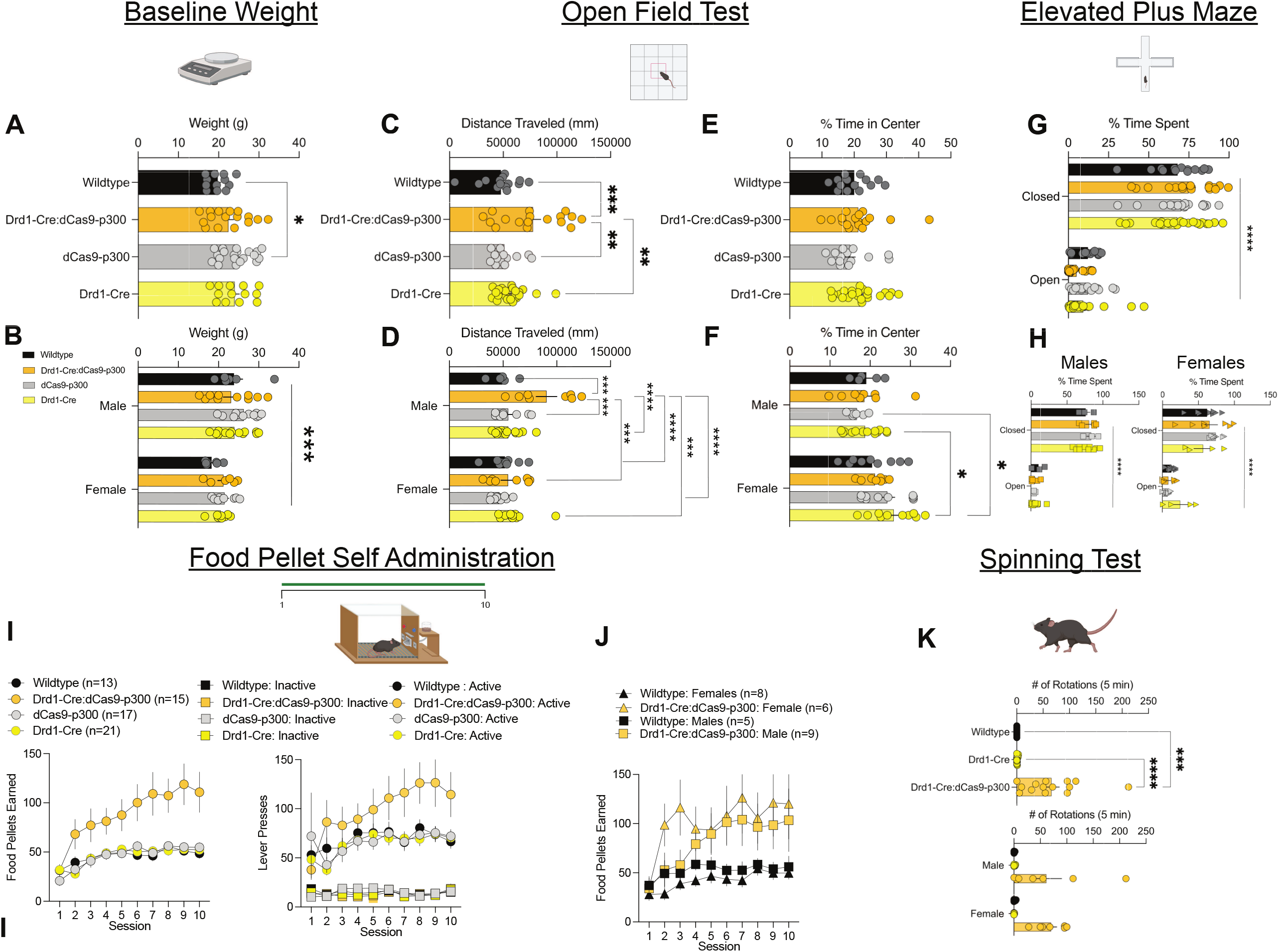
Drd1-Cre:dCas9-p300 mice have augmented baseline locomotion, stereotypic spinning and reward-learning behaviors. **A**, Adult dCas9-p300 mice exhibit increased average weight in comparison to wildtype mice, however when **B**, compared by sex, this effect is suggested to be due to sex and not genotype among the Drd1-Cre x dCas9-p300 crosses. **C**, Drd1-Cre:dCas9-p300 mice display stereotypic spinning behavior in comparison to wildtype and Drd1-Cre mice. **D**, Both male and female Drd1-Cre:dCas9-p300 mice display stereotypic spinning at similar levels. **E**, Drd1-Cre:dCas9-p300 mice have significantly higher locomotor behavior in an open field fest in comparison to wildtype, dCas9-300, and Drd1-Cre mice. **F**, Male Drd1-Cre:dCas9-p300 mice display significantly higher locomotion than female Drd1-Cre:dCas9-p300, as well as all other male and female genotypes. **G**, No changes in anxiety-like behaviors were detected from the open field test between Drd1-Cre:dCas9-p300, dCas9-p300, Drd1-Cre, and wildtype mice. **H**, However, female mice across genotypes spent significantly more time in the center of the open field test in comparison to male mice. **I**, No differences were observed in anxiety-like behaviors from the elevated plus maze across all genotypes. **J**, Both male and female mice of all genotypes spent significantly more time in the closed versus open arm. **K**, Drd1-Cre:dCas9-p300 mice earned significantly more food pellets and made higher number of active lever presses during operant food self-administration than all other genotypes. **L**, Drd1-Cre:dCas9-p300 mice earned more food pellets during operant food self-administration than wild-type mice regardless of sex. **p<0.01, ***p<0.001. Each bar represents mean ±SEM. *p<0.05

#### Baseline Anxiety-like Behavior and Reward-Learning

To determine whether Drd1-specific expression of dCas9-p300 impacts baseline anxiety-like behavior, changes in percent of time spent in the center of field during the open field test was examined. No differences were seen between Drd1-Cre:dCas9-p300, dCas9-p300, Drd1-Cre, and wildtype mice (**Fig. 3E**: Kruskal-Wallis: H(3)=3.3866, p=0.3358). dCas9-p300 and Drd1-Cre female mice spent significantly more time in the center of the open field test in comparison to their respective males (**Fig. 3F**: two-way ANOVA; Interaction: F_(3,61)_=1.52, p=0.2177; Sex: F_(1,61)_=14.4, p=0.0003; Genotype: F_(3,61)_=1.09, p=0.3592; Fishers’s post-hoc: males vs. females dCas9-p300, Drd1-Cre:p<0.01).

Baseline effects of Drd1-dCas9-p300 expression on anxiety-like behavior was further assessed using the elevated plus maze. All genotypes spent significantly more percent of time in closed arms versus open arms (**Fig. 3G:** two-way ANOVA; Interaction: F_(3,61)_=1.254, p=0.2974; Arm: F_(1,65)_=482.7, p<0.0001; Genotype: F_(3,65)_=0.4160, p=0.7421). Both male and female mice spent significantly more time in closed versus open arms (**Fig. 3H**: male: two-way ANOVA; Interaction: F_(3,17)_=0.4385, p=0.7284; Arm: F_(1,17)_=310.9, p=0.0001; Genotype: F_(3,1*7*)_=0.3243,p=0.8078; female: two-way ANOVA; Interaction: F_(3,23)_=1.558, p=0.2266; Arm: F_(1,23)_=86.62, p=0.0001; Genotype: F_(3,23)_=0.4295,p=0.7338). Overall, no baseline differences in anxiety-like behavior were detected between Drd1-Cre:dCas9-p300 mice and their genotype littermates. Given that D1-SPN activity regulates reward learning, the effects of dCas9-p300 expression in Drd1-expressing cells on acquisition of food self-administration were assessed. Drd1-Cre:dCas9-p300 mice earned significantly more food pellets (**Fig. 3I**: two-way ANOVA: Interaction: F_(27,540)_=2.852; p<0.001; Session: F_(1.622, 97.31)_=20.45; p<0.001; Genotype: F_(3,60)_=8.088; p=0.001; Tukey’s *post-hoc*: Drd1-Cre:dCas9-p300 vs. wildtype on Session 8,9,10: p<0.05; Drd1-Cre:dCas9-p300 vs. Drd1-Cre on Session 8,9) or pressed more active, but not inactive lever (two-way ANOVA: Interaction: F_(63, 1053)_=1.917; p<0.01; Session: F_(2.755, 322.3)_=6.091; p<0.001; Genotype and Lever: F_(7,117)_=26.66; p<0.001; Fisher’s LSD *post-hoc*: Session 8: Drd1-Cre:dCas9-p300 vs Drd1-Cre: p<0.05; Drd1-Cre:dCas9-p300 vs dCas9-p300 p<0.05) during food self-administration sessions than Drd1-Cre, dCas9-p300 and wildtype counterparts. No sex specific differences on the amount of food pellets earned during operant food self-administration were observed between dCas9-p300 mice or wildtype counterparts (**Fig. 3J:** three-way ANOVA: Session x Genotype: F(9,216)=0.6826; p=0.7242; Sex x Genotype: F(1,24)=0.8466; p=0.3667; Sex x Genotype x Session: F(9,216)=0.9506; p=0.4822; Session: F(9,216)=7.540; ***p<0.001; Sex: F(1,24)=0.091; p=0.7655; Genotype: F(1,24)=7.281; *p<0.05). Overall, these findings would suggest augmented baseline reward-learning from dCas9-p300 expression in Drd1-cells.

#### Baseline Stereotypic Behavior

To assess stereotypic behavior, mice underwent a spinning test. Drd1-Cre:dCas9-p300 mice displayed robust spinning stereotypy in comparison to wildtype and Drd1-Cre mice (**Fig. 3K:** one-way ANOVA; F_(2,31)_=13.38, p<0.001; Tukey’s post-hoc: wildtype vs Drd1-Cre:dCas9-p300: p<0.001; Drd1-Cre vs Drd1-Cre:dCas9-p300: p<0.001) Stereotypy was observed both male and female Drd1-Cre:dCas9-p300 mice, with stereotypic spinning at similar levels (**Fig. 3K:** two-way ANOVA; Interaction: F_(2,32)_=0.07, p=0.9281; Sex: F_(1,32)_=0.06; p=0.8030; Genotype: F_(2,32)_=14.7, p<0.001; Fisher’s *post-hoc:* males: wildtype vs Drd1-Cre:dCas9-p300: p<0.001; Drd1-Cre vs Drd1-Cre:dCas9-p300: p<0.001; females: wildtype vs Drd1-Cre:dCas9-p300: p<0.001; Drd1-Cre vs Drd1-Cre:dCas9-p300: p<0.001).

### Ador2a-Cre:dCas9-p300 mice display typical baseline behaviors

#### Adult Baseline Weight and General Activity

The effects of Cas9-p300 expression in Ador2a-cells on bodyweight and home-cage behaviors were also assessed on Ador2a-Cre:dCas9-p300 mice and their littermates. No differences in adult baseline weights were seen across genotypes (**Fig. 4A:** one-way ANOVA; F(3,45)=1.890, p=0.1488). As expected, male mice had increased weight across all genotypes (**Fig. 4B:** Two-way ANOVA: Interaction: F(3,41)=0.9326 p=0.4337; Sex: F(1,41)=15.76, ***p<0.001; Genotype: F(3,41) 0.3052, p=0.8214). No notable differences in grooming, appearance or activity were observed in the homecage. Examinations of baseline locomotion in open field testing confirmed no differences in locomotion across genotype (**Fig. 4C**: one-way ANOVA: F(3, 53)=1.442, p=0.2140) or sex (**Fig. 4D**: two-way ANOVA; Interaction: F(3,52)=1.37, p=0.2614; Sex: F(1,52)=1.87, p=0.1768; Genotype: F(3,52)=0.09,p=0.9599). Overall, these data would suggest that dCas9-p300 expression in Ador2a cells have no impact on general activity and development in mice.

**Figure 4.**
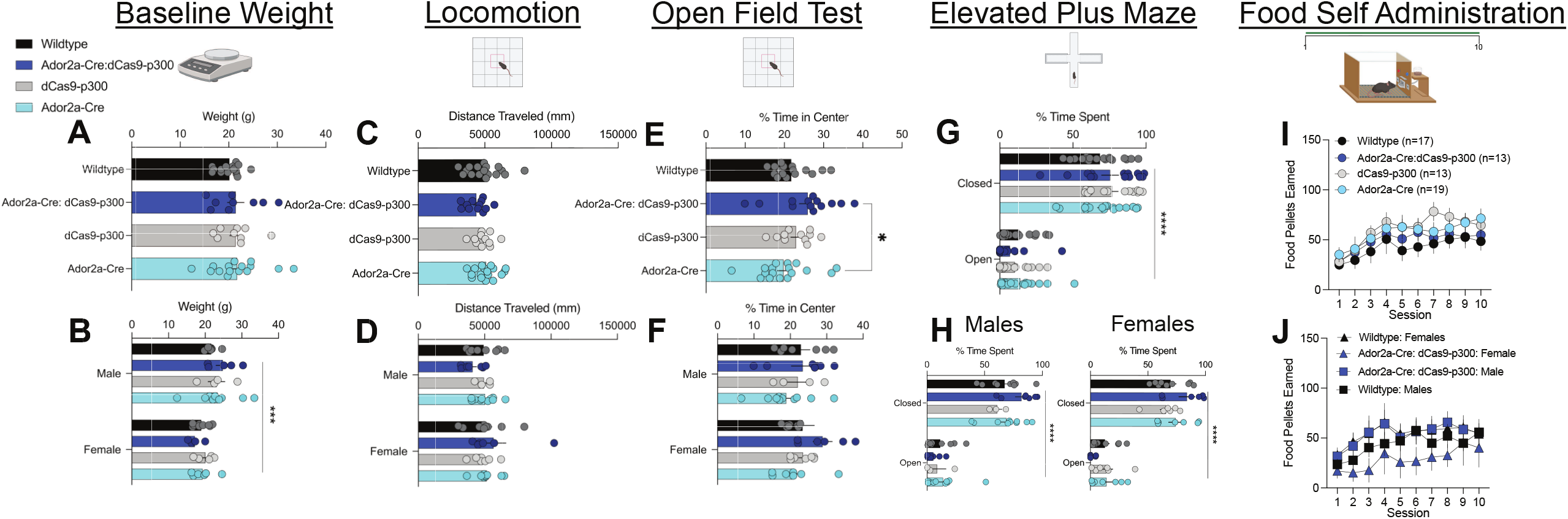
Ador2a-Cre:dCas9-p300 mice display typical baseline behaviors. **A**, Adult wildtype, Ador2a-Cre:dCas9-p300, dCas9-300 and Ador2a-Cre mice have similar average weights, with **B**, male mice showing increased weight across all genotypes. **C,D** No differences in locomotion were seen across genotype or sex. **E,F** Ador2a-Cre:dCas9-p300 mice spent higher percentage of time in the center of the box during the open field test in comparison to Ador2a-Cre mice. However, no sex differences were seen in percent time spent in the center. **G,H**, No differences were detected from time spent between arms in the elevated plus maze across all genotypes. Male and female mice of all genotypes also spent significantly more time in the closed versus open arm. **I**, No differences in food pellets earned were seen across all genotypes in operant food self-administration. **J**, Ador2a-Cre:dCas9-p300 male and female mice earned similar amount of food pellets during operant food self-administration as wild-type male and female mice. Each bar represents mean ±SEM. *p<0.05, **p<0.01, ***p<0.001.

#### Baseline Anxiety-like Behavior and Reward-Learning

The effects of Ador2a-specific expression of dCas9-p300 on baseline anxiety-like behavior were assessed using percent of time spent in center of field during the open field test. Ador2a-Cre:dCas9-p300 spent more percent of time in the center of the open field test in comparison to Ador2a-Cre mice (**Fig. 4E:** one-way ANOVA; F(3,53)=0.6579, p=0.0531 Tukey’s test: Ador2a-Cre:dCas9-p300 vs Ador2a-Cre: p<0.05). However, no sex specific differences were observed (**Fig. 4F:** two-way ANOVA: Interaction: F(3,49)=0.31, p=0.8134; Sex: F(1,49)=1.49, p=0.2281; Genotype: F(3,49)=1.89,p=0.1434). Similarly, no differences were detected in time spent between arms in the elevated plus maze (**Fig. 4G:** two-way ANOVA: Interaction: F(3,59)=0.9856, p=0.4058; Arm: F(1,59)=330.3, p<0.0001; Genotype: F(3,59)=0.6883,p=0.5628). Male mice of all genotypes did spend significantly more time in the closed versus open arm (**Fig. 4H:** two-way ANOVA; Interaction: F(3,25)=0.1.34, p=0.2820; Arm: F(1,25)=135, p<0.0001; Genotype: F(3,25)=1.33,p=0.2545) and female (**Fig. 4H:** two-way ANOVA; Interaction: F(3,24)=0.181, p=0.1707; Arm: F(1,24)=175, p<0.0001; Genotype: F(3,24)=1.306,p=0.3825). Altogether, these results indicate dCas9-p300 expression within Ador2a-cells in mice has no impact on anxiety-like behaviors.

Given the known roles of A2A-SPNs on reward-learning and potential application of Ador2a-Cre:dCas9-p300 mice in reward-based tasks, genotypic differences on acquisition of food self-administration were measured. Unlike Drd1-Cre:dCas9-p300 mice, no differences in food pellets earned (**Fig. 4I**; two-way ANOVA; Interaction: F_(27,477)_=1.053, p=0.3942; Session: F_(1.675, 88.78)_=22.52, p<0.0001; Genotype: F_(3,53)_=0.8858,p=0.4545) were seen across all genotypes during operant food self-administration. Similarly, both Ador2a-Cre:dCas9-p300 male and female mice earned similar amount of food pellets during operant food self-administration as wild-type male and female mice (**Fig. 4J**; two-way ANOVA; Session: F_(9,220)_=3.108, p<0.00015; Sex: F_(1,220)_=2.798e-005, p=0.9958, Genotype: F_(1,220)_=3,393,p=0.0668; Sex x Genotype: F_(1,220)_=8.379,p=0.0042). These results suggest that expression of dCas9-p300 within Ador2a-cells in mice has no impact on reward learning.

### Drd1-expressing cells in Drd1-Cre:dCas9-p300 mice have enhanced sensitivity to synaptic input

We hypothesized that dCas9-p300 expression enhances D1-SPN activity, given that activation of D1-SPNs promotes locomotor and motivation-based behaviors, as similarly seen in Drd1-Cre:dCas9-p300 mice (Lobo et al., 2010; Lee et al., 2018). To determine the effects of dCas9-p300, whole-cell patch clamp recordings were performed in Drd1-expressing cells within the DS of Drd1-Cre:dCas9-p300 and Drd1-Cre mice. To visualize D1-SPNs, AAV9-hSyn-DIO-mCherry was infused in the DS of Drd1-Cre:dCas9-p300 or Drd1-Cre mice. Whole-cell recordings were performed in current-clamp mode and baseline membrane properties of mCherry positive (mCherry+) cells were observed (**Fig 5A**.). There was no significant difference in the number of action potentials at increasing current steps between the two genotypes (**Fig 5B**.). In addition, Drd1-Cre:dCas9-p300 showed no difference in rheobase or threshold potential compared to Drd1-Cre mice. However, Drd1-Cre:dCas9-p300 mice had significantly higher input resistance than Drd1-Cre mice (**Fig 6C**.: un-paired t-test: t(15)=2.45; p=0.0270). There was a significant increase in sEPSC amplitude of mCherry+ cells within Drd1-Cre:dCas9-p300 compared to Drd1-Cre mice (**Fig 5D**.: un-paired t-test: t(9.562)=2.387, p=0.0392) but no significant difference in inter-event intervals (un-paired t-test: t(14)=0.8044, p=0.4346) were observed. Altogether, these results suggest that Drd1-Cre:dCas9-p300 mice have increased glutamatergic synaptic drive of D1-SPNs.

**Figure 5.**
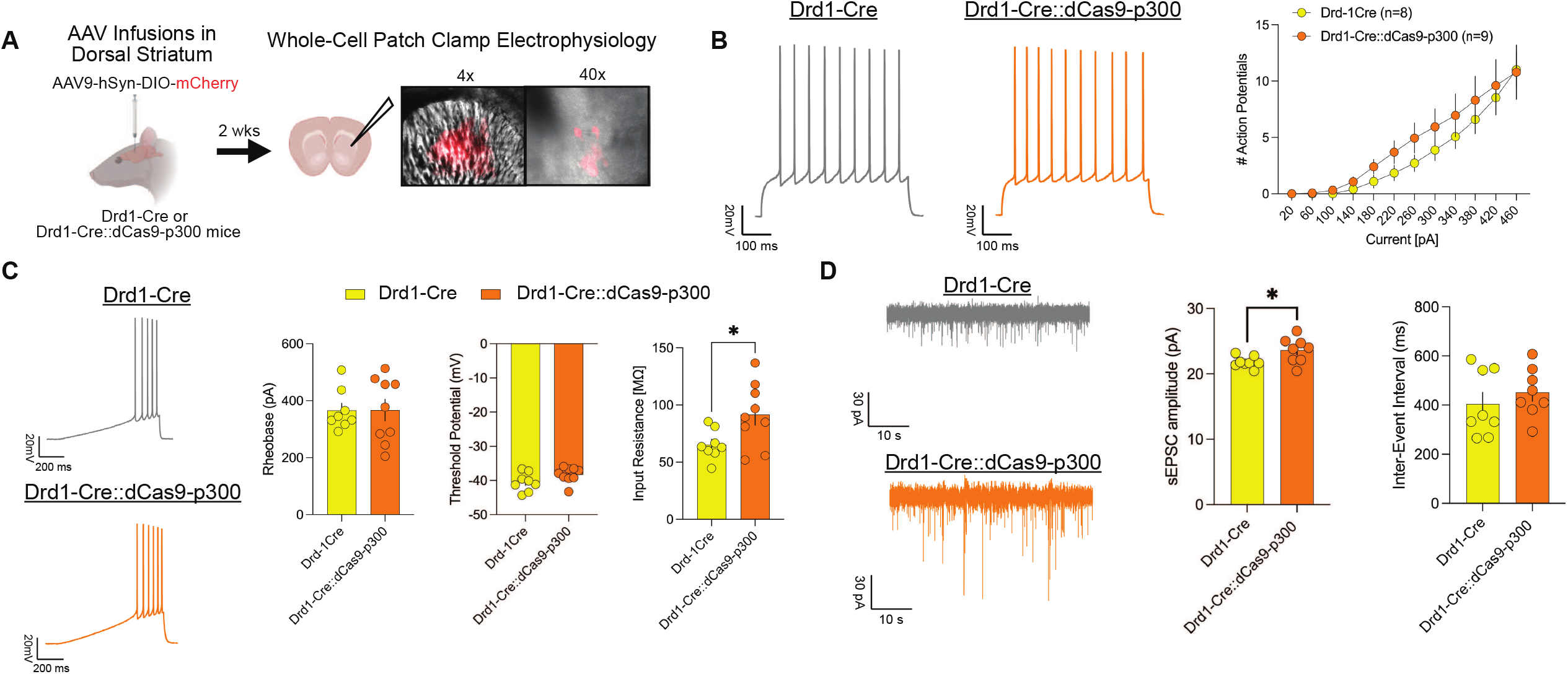
dCas9-p300 expression in dorsal striatal D1-SPNs enhances excitatory drive in these neurons. **A**, Drd1-Cre and Drd1-Cre:dCas9-p300 mice underwent bilateral infusions of AAV9-DIO-hSyn-mCherry in the DSt for whole-cell patch clamp electrophysiology. **B**, mCherry-positive cells recorded from Drd1-Cre:dCas9-p300 mice exhibit no difference in action potential (AP) number in comparison to Drd1-Cre mice. **C**, mCherry-positive cells from Drd1-Cre:dCas9-p300 mice exhibit no difference in rheobase or threshold potential compared to Drd1-Cre mice. Input resistance was significantly higher in Drd1-Cre:dCas9-p300 mCherry-positive cells compared to Drd1-Cre mice, despite no detected changes in membrane capacitance or resting membrane potential between genotypes **D**, There is significantly higher amplitude of spontaneous excitatory postsynaptic current (sEPSC) events in Drd1-Cre:dCas9-p300 mCherry-positive cells compared to Drd1-Cre mice, but no difference in sEPSC inter-event interval. Each graphed bar and line represent mean ± SEM

## Discussion

Tools that employ Cre-dependent dCas9 and epigenome modifier fusions are being developed for basic research to perform cell-type specific manipulation of gene expression. Cre-dependent transgenic mouse lines expressing dCas9 epigenome editors are being generated to circumvent the technical difficulties from using CRISPR-based viral vector, and simply allowing for viral delivery of sgRNAs alone. Here, we characterized the impacts of expressing dCas9-p300 in Drd1 or Ador2a cell types, as the first report of breeding LSL-dCas9-p300 with Cre-driver mouse lines. We demonstrate that expression of dCas9-p300 in Drd1 expressing cells causes altered locomotor and reward-based behaviors at baseline, along with enhanced sensitivity to synaptic input in D1-SPNs. However, we found no observable effects on behavior from dCas9-p300 in Ador2a expressing cells. Altogether, this would suggest cautionary experimental approach for pairing LSL-dCas9-p300 lines with Cre-driver mouse lines for neuroscience research, particularly for behavioral studies.

Our data demonstrates that pairing LSL-dCas9-p300 mice with Cre-driver mouse lines leads to Cre-dependent expression of dCas9-p300 that is predominantly cell-type specific. Because the p300 core fused to dCas9 is a constitutively active catalytic domain of the human p300 acetyltransferase, it was critical to know whether its ambient expression creates unintentional alterations of gene expression in our Drd1-Cre- or Ador2a-Cre:dCas9-p300 lines. However, we found minimal effects of dCas9-p300 on baseline striatal expression of genes related to the transgene expression of either Drd1-Cre- or Ador2a-Cre:dCas9-p300 lines. Although DSt *p300* expression within Drd1-Cre:dCas9-p300 was elevated in comparison to dCas9-p300, this pattern was absent in Ador2a-Cre:dCas9-p300 lines. Within Ador2a-Cre:dCas9-p300 lines, DSt *Ador2a* was reduced in comparison to wildtype mice, which may suggest counter-adaptive effects of Ador2a expression due to the dCas9-p300 transgene. Widespread changes in both H3K27ac levels and gene expression were discovered from ChIP- and RNA-Seq within the liver of dCas9-p300 mice that were IV injected with AAV-Cre and Pdx1-targeted sgRNAs in comparison to saline infused controls in the original report (Gemberling et al., 2021). It is plausible that off-target histone acetylation and gene expression changes are present within our generated mouse lines, despite not detecting similar sgRNA independent consequences from the expression of dCas9-p300 alone. Additionally, p300 directly binds to other transcriptional machinery proteins, therefore it is possible that dCas9-p300 sequesters these complexes away from DNA to indirectly impact transcription (Gerritsen et al., 1997; Hóbor et al., 2022). Future characterization studies on dCas9-p300 within Cre-driver mouse lines would therefore benefit from utilizing cell-type specific RNA isolation methods.

Drd1-Cre:dCas9-p300 mice show stereotypic, spinning behavior, hyperlocomotion and enhanced acquisition of reward tasks. These behaviors are often seen following activation of Drd1-expressing cells within the striatum, suggesting Drd1-Cre:dCas9-p300 mice may have hyperactive Drd1 cellular activity. Indeed, we observed that Drd1-Cre:dCas9-p300 D1-SPNs have enhanced excitatory synaptic input as well as a possible elevated sensitivity to this input in the form of increased input resistance. The detected increases in input resistance and sEPSC amplitude may reflect a decrease in the number, or conductance, of inward rectifying potassium or slowly inactivating A-type potassium channels that, depending on SPN voltage state, may enhance excitatory synaptic drive (Wilson, 2009; Steephen and Machanda, 2009; John and Manchanda, 2011). Our previous study investigating a Drd1 specific knockout of tropomysin receptor kinase B (TrkB), the receptor for brain derived neurotrophic factor (BDNF), Drd1-Cre::flTrkB, demonstrated that a subset of this genotype displayed stereotypy behavior including enhanced spinning similar to Drd1-Cre:dCas9-p300 mice (Engeln et al., 2021). The Drd1-Cre::flTrkB stereotypy mice had reduced dendritic complexity in striatal D1-SPNs compared to wildtype controls or Drd1-Cre::flTrkB non-stereotypy mice. It is plausible that Drd1-Cre:dCas9-p300 D1-SPNs also display altered dendritic complexity that would further affect dendritic integration and, ultimately,action potential generation. Given that p300 is a transcriptional activator dCas90-p300 expression may promote expression of genes that regulate intrinsic factors related to ion channel activity and dendritic complexity. This line can further be studied to parse out underlying mechanisms of stereotypy behavior and reward-learning (Engeln et al., 2021). Future studies are positioned to examine overlapping mechanisms between Drd1-Cre:dCas9-p300 and Drd1-Cre::flTrkB stereotypy lines to understand why the observed morphological and physiological changes in D1-SPNs result in this behavioral phenotype.

Although activation of A2A-SPNs generally reduces locomotion, Ador2a-Cre:dCas9-p300 have no changes in locomotion across other genotypes (Kravitz et al., 2010; Zhu et al., 2016). Similarly, activation of A2A-SPNs can promote negative affective behaviors in mice, yet we report no effects on anxiety-like behaviors were seen in Ador2a-Cre:dCas9-p300 mice (Francis et al., 2015; Correia et al., 2023). This may suggest different downstream effects of histone acetylation and gene expression related to A2A-SPN neural activity and behavior in comparison to R-SPN activity and behavior. In support of this, disruption of the Class I histone deacetylase HDAC3’s activity in D1-SPNs but not A2A-SPNs impacts reward-related behaviors such as cocaine-seeking (Campbell et al., 2021).

Alternatively, dCas9-p300 may indeed result in activation of A2A-SPNs, however compensation of A2A-SPN activity may have occurred to prevent behavioral deficits. Furthermore, temporal specificity of A2A-SPN activity may play a key role in causing behavioral changes. This is seen from studies showing that activation of Ador2a-expressing cells can enhance or diminish motivated behaviors when engaged at different time points in reinforcement-related tasks (Soares-Cunha et al., 2016, 2022; Cole et al., 2018). Ultimately, should this line be pursued then follow-up studies examining whether the cellular physiology of the A2A-SPN cell types is altered from dCas9-p300 expression are imperative in interpreting these behavioral results.

It is imperative to determine the limitations of new biological tools prior to their implementation into disease-based research endeavors. Although the LSL-dCas9-p300 shows promising applications for neuroscience studies, our data outlines reasons to proceed with caution when crossing this mouse line with Cre driver lines, particularly for behavior related studies. It plausible that the disrupted behavior and D1-SPN physiology observed in the Drd1-Cre:dCas9-p300 mice is a consequence of developmental expression of the transgene. Further testing with inducible Cre systems or viral Cre in adulthood should be examined in the dCas9-p300 mouse line. For future directions, it is possible that alternative CRISPR-related transgenic mouse lines using other epigenomic editors fused to dCas9 lead to fewer off target effects, including activators such as VP64 or VPR or repressors such as KRAB or HDACs. dCas9-p300 expression throughout development may have attributed to these behavioral disruptions, as there are no reported off-target effects with viral based approaches using dCas9-p300 in the brain. Overall, our current findings suggest that investigators should characterize the impacts of LSL-dCas9-p300 in Cre driver mouse lines prior to the introduction of sgRNAs in behavior-related studies.

## Acknowledgments

This work was supported by R21/R33DA05210, R01DA038613, R01DA047843 and to M.K.L; F32DA056191 to R.R.C., F31DA052967 to E.Y.C., and F99NS135696 to D.F.

## Author Contributions

RRC, EYC and MKL designed the study. RRC, MG, ABW, AS and MKL wrote the manuscript. RRC, EYC, MG, ABW, and AS analyzed the data. RC, EYC, MG, CB and SK performed all behavioral experiments. RRC, MG, and SK performed stereotaxic surgeries for electrophysiological experiments and ABW and AS performed electrophysiological recordings. RRC, EYC, MG, GLV, SK, and SP performed RT-qPCR including tissue collection, RNA isolation and RT-qPCR. RRC, MG, GLV, SP, SSK, performed RNAScope experiments, ranging from tissue preparation, microscopy and image analysis. MG performed western blotting. RRC, EYC, SK, and MB helped maintain transgenic mouse lines and oversee colony maintenance. DF and SG assisted with data storage and figure preparation.

