## Supplemental Figures & Table for "Dopamine receptor 1 specific CRISPRa mice exhibit disrupted behaviors and striatal baseline cellular activity"

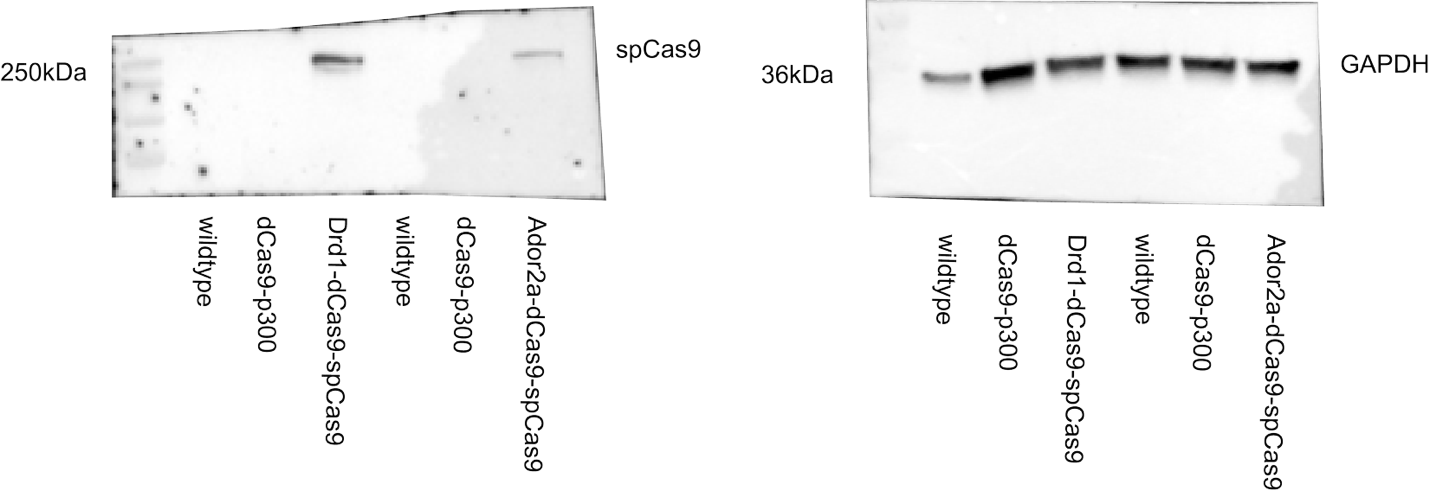

Figure S1. Full Western blot data of CRISPRi mice for Cas9 expression validation. Samples in order from left to right are: wildtype, dCas9-p300, Drd1-Cre:dCas9-p300, wildtype, dCas9-p300 and Adora2a-Cre:dCas9-p300. The blot was cut at the 50kDa ladder mark. On the left is the half of the blot that was >50kDa which was probed with an Anti-Cas9 antibody. Cas9 was detected near the 250kDa marker in only the Drd1-Cre:dCas9-p300 and Adora2a-Cre:dCas9-p300 samples. On the right is the half of the blot that was <50kDa, which was probed with an Anti-GAPDH antibody. GAPDH was detected in all samples.

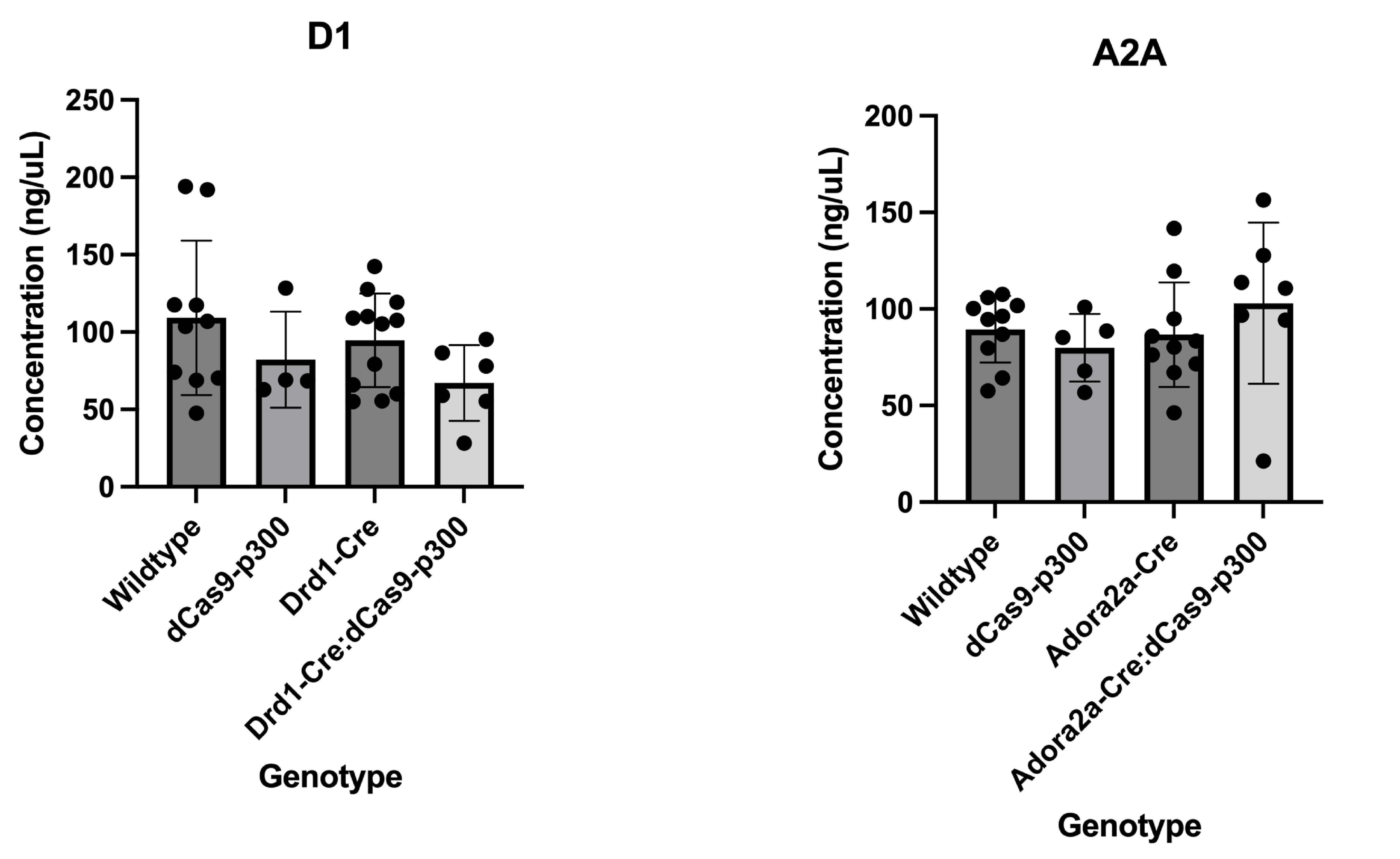

Figure S2. RNA concentration (in ng/µL) from NAc tissue from mice. There was no significant difference in RNA yield between the genotypes for each line.

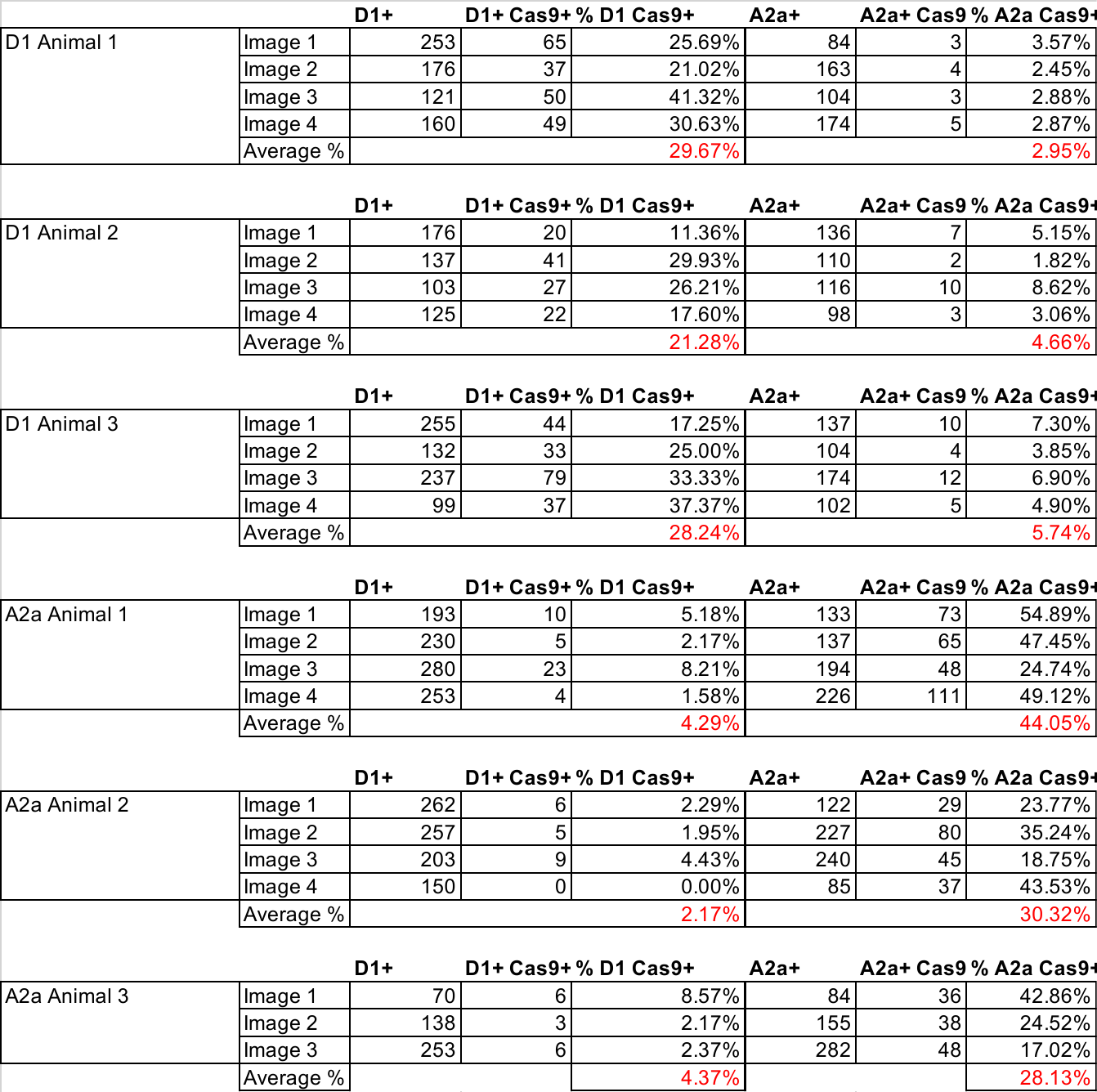

Table S1. RNAscope quantification
